# Dynamics of pellet fragmentation and aggregation in liquid-grown cultures of *Streptomyces lividans*

**DOI:** 10.1101/270595

**Authors:** Boris Zacchetti, Paul Smits, Dennis Claessen

## Abstract

Streptomycetes are extensively used for the production of valuable products, including various antibiotics and industrial enzymes. The preferred way to grow these bacteria in industrial settings is in large-scale fermenters. Growth of streptomycetes under these conditions is characterized by the formation of complex mycelial particles, called pellets. While the process of pellet formation is well characterized, little is known about their disintegration. Here, we use a qualitative and quantitative approach to show that pellet fragmentation in *Streptomyces lividans* is initiated when cultures enter the stationary phase, which coincides with a remarkable change in pellet architecture. Unlike young pellets, aging pellets have a less dense appearance and are characterized by the appearance of filaments protruding from their outer edges. These morphological changes are accompanied by a dramatic increase in the number of mycelial fragments in the culture broth. In the presence of fresh nutrients, these fragments are able to aggregate with other small fragments, but not with disintegrating pellets, to form new mycelial particles. Altogether, our work indicates that fragmentation might represent an escape mechanism from the environmental stress caused by nutrient scarcity, with striking similarities to the disassembly of bacterial biofilms.

## Introduction

Streptomycetes are sporogenic, soil-dwelling bacteria that belong to the phylum *Actinobacteria*. Unlike unicellular bacteria, streptomycetes form so-called mycelia, which are intricate networks of interconnected filamentous cells. As producers of a plethora of valuable secondary metabolites (e.g. antibiotics) and hydrolytic enzymes, streptomycetes are extensively used in the field of industrial biotechnology (Vrancken and Anné, 2009;Barka et al., 2016). For their industrial exploitation, streptomycetes are typically grown in large-scale bioreactors, where they display morphologies that not only differ between different strains, but also greatly depend on the growth conditions (Tresner et al., 1967;Braun and Vecht-Lifshitz, 1991;van Dissel et al., 2014). Typically, morphologies vary from small fragments to large mycelial particles with a diameter of several hundred microns. For instance, the model strain *Streptomyces lividans*, an industrial workhorse for enzyme production, forms dense mycelial structures called pellets. Previous research showed that the size of these pellets is highly heterogeneous (van Veluw et al., 2012), which poses constraints in terms of industrial exploitability. An important factor driving the size heterogeneity of pellets is the stochastic aggregation of germinating spores, which thereby lose their individuality (Zacchetti et al., 2016). Aggregation is mediated by at least two glycans associated with the outer cellular surface. These glycans are produced under control of the *cslA/glxA* operon and the *mat* cluster (Xu et al., 2008;de Jong et al., 2009;Chaplin et al., 2015;van Dissel et al., 2015;Petrus et al., 2016). Whereas the structure of the glycan produced by CslA and GlxA has not yet been resolved, that produced by the Mat proteins was recently shown to be poly-β-(1,6)-Nacetylglucosamine (PNAG) (van Dissel et al., 2018). PNAG is not uniquely present in streptomycetes, but is a commonly identified constituent of the extracellular matrix of bacterial biofilms, where this polymer contributes to the adherence of cells to one another (Jefferson, 2009;Pace et al., 2009).

In addition to genetic factors, environmental factors and culture set-up also influence the growth and morphology of streptomycetes (van Dissel et al., 2014). One poorly-studied process affecting the morphology of mycelia is that of fragmentation. Fragmentation consists in the detachment of mycelial portions from pre-existing particles, an occurrence which has been observed in streptomycetes after prolonged periods of growth in fermenters or under temperature induced stress (Kim and Hancock, 2000;Manteca et al., 2010;Rioseras et al., 2013). Notably, very little is known about the timing of this process in relation to growth and morphology, the fate of the detached fragments, and the role that fragmentation has on culture heterogeneity.

Here, we systematically analyzed the dynamics of fragmentation using a qualitative and quantitative approach. We first show that the architecture of pellets changes when cultures enter the stationary phase. This morphological switch coincides with the detachment of small viable fragments that we show are able to aggregate with one-another and to form new pellets in presence of fresh nutrients. These results not only enhance our understanding of pellet morphogenesis and growth dynamics in *Streptomyces*, but also show that fragmentation is strikingly similar to biofilm dispersal.

## Materials and Methods

### Bacterial strains and culture conditions

The strains used in this study are listed in Table 1. Strains were grown on MS agar medium (Kieser et al., 2000) at 30°C to prepare spore solutions and to quantify the amount of colony forming units in liquid-grown cultures. Liquid cultivations were performed in a total end volume of 100 ml (thus seed culture volumes of 2 or 10 ml were added to 98 and 90 ml fresh medium, respectively). The used liquid media were TSBS (Kieser et al., 2000) or ½ x TSBS medium (15 g/l tryptic soya broth, 10% sucrose). Cultures were grown in 250 ml unbaffled Erlenmeyer flasks equipped with metal coils, unless differently stated. Cultures were grown at 30°C in an orbital shaker with a 2-inch stroke at 160 rpm. All seed cultures were inoculated with 10^6^ spores ml^-1^. For co-cultivations, 5 x 10^5^ spores ml^-1^ of each fluorescent strain were used. Dry weight measurements were performed with the freeze-dried mycelium obtained from 10 ml culture broth, after having washed the biomass twice with 10 ml milliQ water.

**Table 1.** *Streptomyces* strains used in this study

| Strain | Description | Reference |
| --- | --- | --- |
| *Streptomyces lividans* 66 | Wild-type strain | Laboratory stock |
| *Streptomyces lividans* pGreen | *S. lividans* 66 containing pGreen | Zacchetti *et al.* 2016 |
| *Streptomyces lividans* pRed* | *S. lividans* 66 containing pRed* | This study |

### Filtration and analysis of seed cultures

Seed cultures were grown for 24, 48 and 72 h before being sequentially filtered through 100, 40 and 5 µm filters (Falcon^®^ Cell Strainer 100 µm Nylon, Falcon^®^ Cell Strainer 40 µm Nylon, PluriSelect pluriStrainer^®^ 5 µm). The 40 µm filtration step was necessary to prevent clogging of the 5 µm filter. The 48- and 72-h samples were inspected without further preparation, whereas for the 24-h sample 15 ml of the filtrate were concentrated via centrifugation at 1000 rpm for 30 min at 4° C and subsequently analyzed.

### Quantification of fragmentation

Large aggregates and pellets present in 50 ml seed cultures at 16, 24, 32 or 40 h, were filtered using a 100 µm filter (Falcon^®^ Cell Strainer 100 µm Nylon). Small particles potentially entrapped in the filtered biomass were removed by four consecutive washing steps, for which 25 ml sterile milliQ water were allowed to pass through the filter. The filtered aggregates and pellets were subsequently brought into sterile Erlenmeyer flasks by rinsing the filter bottom up with 100 ml spent medium, which was obtained by filter-sterilizing the original culture medium using a filtration device equipped with a 0.22 µm filter (Sarstedt Filtropur^®^ V50 500 ml). The cultures were then grown for another 8 h, after which they were again filtered through a 100 µm filter. At this stage pellets were discarded, while the filtrate was used to quantify the number of Colony Forming Units (CFUs, see below), or to inoculate new TSBS cultures, for which 50 ml filtrates were mixed with 50 ml fresh TSBS medium.

### Quantification of Colony Forming Units (CFUs) in liquid-grown cultures

Samples of liquid-grown cultures or filtrates thereof were diluted in MQ water before being plated on MS agar plates. To this end, 1 ml of the diluted samples was evenly distributed on the surface by gently swirling the plates, after which these were allowed to dry in the fume hood for 30 mins. Colonies were counted after 48 h of growth at 30°C.

### Construction of the reporter plasmid pRed*

To create pRed*, the *gap1* (SCO1947) promoter of *Streptomyces coelicolor* A3(2) M145 and the *mCherry* gene were amplified by PCR with the Gap1-FW primer (GAT<u>AGATCT</u>CCGAGGGCTTCGAGACC) and the mCherry-RV primer (TTTG<u>CGGCCG</u>CTTACTTGTACAGCTCGTCCAT). A pIJ2925-derivative carrying these elements was used as a template (Zacchetti et al., 2016). The obtained PCR product was then ligated as a *Bgl*II-*Not*I fragment into pIJ8630 (Sun et al., 1999). The obtained reporter construct was introduced in *S. lividans* 66 via conjugation (Kieser et al., 2000).

### Microscopy settings

Fluorescence and light microscopy, including whole-slide image analysis were performed as described previously (Zacchetti et al., 2016;Willemse et al., 2017). At least 300 pellets were analyzed for each sample, both to determine Feret diameters as well as the percentages of aggregated pellets in co-cultures of fluorescent strains. For the visualization of viable and dead mycelium, samples were stained with Syto-9 and propidium iodide (PI) (Invitrogen). To this end, Syto-9 and PI were added to the samples prior to imaging at a final concentration of 5 µM and 15 µM, respectively. Stained samples were excited at 488 and 543 nm to detect Syto-9 and PI, respectively. The fluorescence emission of Syto-9 was monitored in the region between 505-545 nm, while a long-pass filter at 560 nm was used to detect PI. The fluorescence micrographs presented in Fig. S5 were background corrected by setting the fluorescence signal outside spores and hyphae to 0, as to obtain a sufficiently dark background. These corrections, as well as all image-analysis based quantifications, were performed using ImageJ version 1.48f.

## Results

### Qualitative and quantitative analysis of fragmentation in *Streptomyces lividans*

With the aim to investigate the onset of fragmentation in liquid-grown cultures, *Streptomyces lividans* 66 was grown in seed cultures for 24, 48 or 72 h. We then used 2 or 10 ml of these seed cultures to inoculate new TSBS cultures, which were subsequently grown for 24 h before being analyzed. Microscopy revealed that transferring mycelium from seed cultures after 24 h yielded pellets with an average Feret diameter of 533 and 483 µm for inoculated volumes of 2 and 10 ml, respectively (Fig. 1, Fig. S1, Table 2). These pellets were significantly larger than those formed after 48 h in the seed cultures, and which had not been transferred to fresh medium. When 48- or 72-h old seed cultures were used to inoculate new cultures, two populations of pellets with distinct sizes became evident after 24 h of growth; in addition to large pellets (with an average diameter larger than 350 µm), many small pellets with a diameter below 250 µm were found. Notably, the size of small and large pellets in the diluted cultures was reduced when a larger seed culture volume was used as the inoculum, independent of its age (Fig.1, Fig. S1, Table 2).

**Figure 1.**
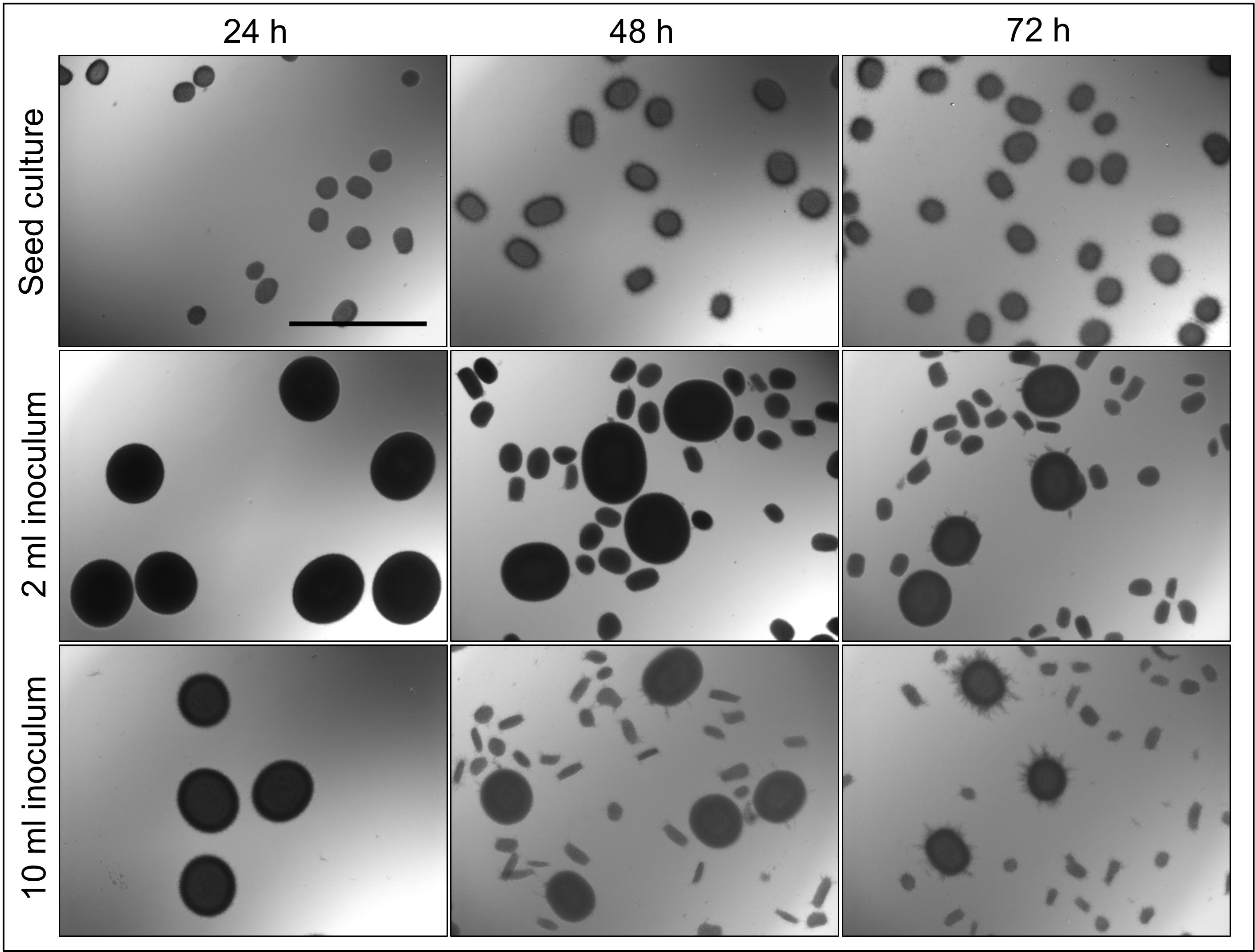
Morphological analysis of pellets of *Streptomyces lividans* 66 in TSBS cultures. The morphology of pellets in seed cultures after 24 (left), 48 (middle) and 72 (right) h of growth in TSBS medium is shown in the top panels. The other panels show micrographs of pellets following the transfer of 2 (middle panels) or 10 ml (bottom panels) of seed cultures in fresh TSBS medium and subsequent growth for 24 h. The scale bar represents 1 mm.

**Table 2.** Average Feret diameter of pellets in 24-hours old liquid cultures, which had been inoculated with 2 or 10 ml of seed cultures of different ages. The average Feret diameter of pellets in the seed cultures is shown inside parentheses.

| Seed culture age<br>(Feret diameter in $\mu\text{m}$) | Transferred seed<br>culture volume | Feret diameter<br>in $\mu\text{m}$ |
| --- | --- | --- |
| 24 h ($194 \pm 28.3$) | 2 ml | $533 \pm 53.4$ |
| | 10 ml | $483 \pm 55.2$ |
| 48 h ($278 \pm 42.1$) | 2 ml | $250 \pm 102$ |
| | 10 ml | $216 \pm 100$ |
| 72 h ($233 \pm 28.3$) | 2 ml | $184 \pm 79.7$ |
| | 10 ml | $154 \pm 78.6$ |

The appearance of small pellets when using seed cultures that were 48- or 72-h old prompted us to investigate whether these originated as a result of the detachment of mycelial fragments from pre-existing pellets in the seed cultures. This hypothesis was reinforced by the observation that the outer edge of pellets after 48 or 72 h of growth was visibly less compact than that of pellets that had grown for 24 h (Fig. 2, Fig. S2). In particular, many filaments were found to protrude from the outer edge of older pellets. To shed light on this, we collected the filtrates of seed cultures after passage through a series of filters, of which the last one had a diameter of 5 µm (see Material and Methods). Interestingly, filtrates of seed cultures grown for 24 h contained approximately the same number of CFUs in the 100 and 5 µm filtrates (1.67*10^5^ ml^-1^ and 1.62*10^5^ ml^-1^ respectively, Table 3). This indicates that the vast majority of particles in the filtrates at this stage was smaller than 5 µm. A microscopic analysis revealed that the large majority of these CFUs were viable ungerminated spores, which had been used for inoculation (Fig. 3). This assumption is based on the fact that *S. lividans* has never been observed to sporulate in TSBS cultures.

**Figure 2.**
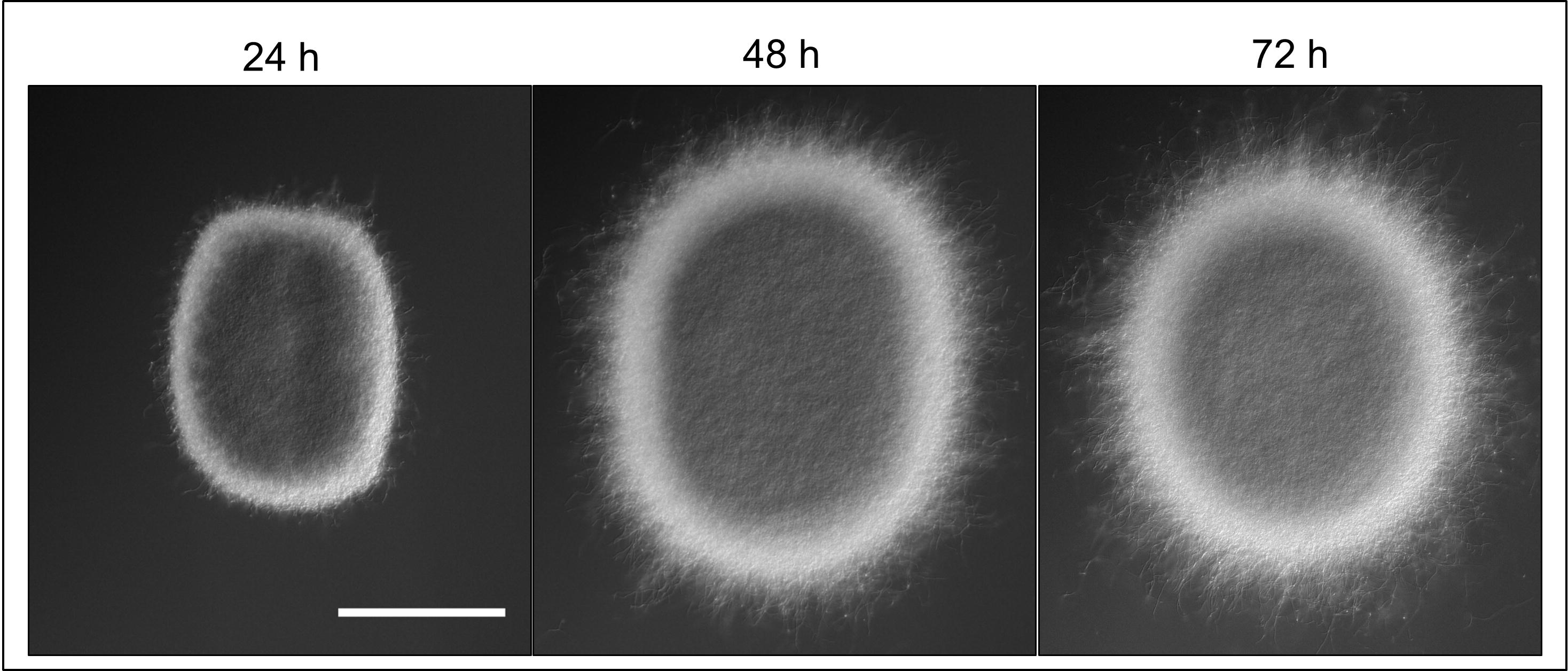
Morphological changes of aging *S. lividans* pellets. Representative pellets of *Streptomyces lividans* 66 after 24, 48 and 72 h of growth in TSBS cultures. The scale bar represents 100 µm.

**Figure 3.**
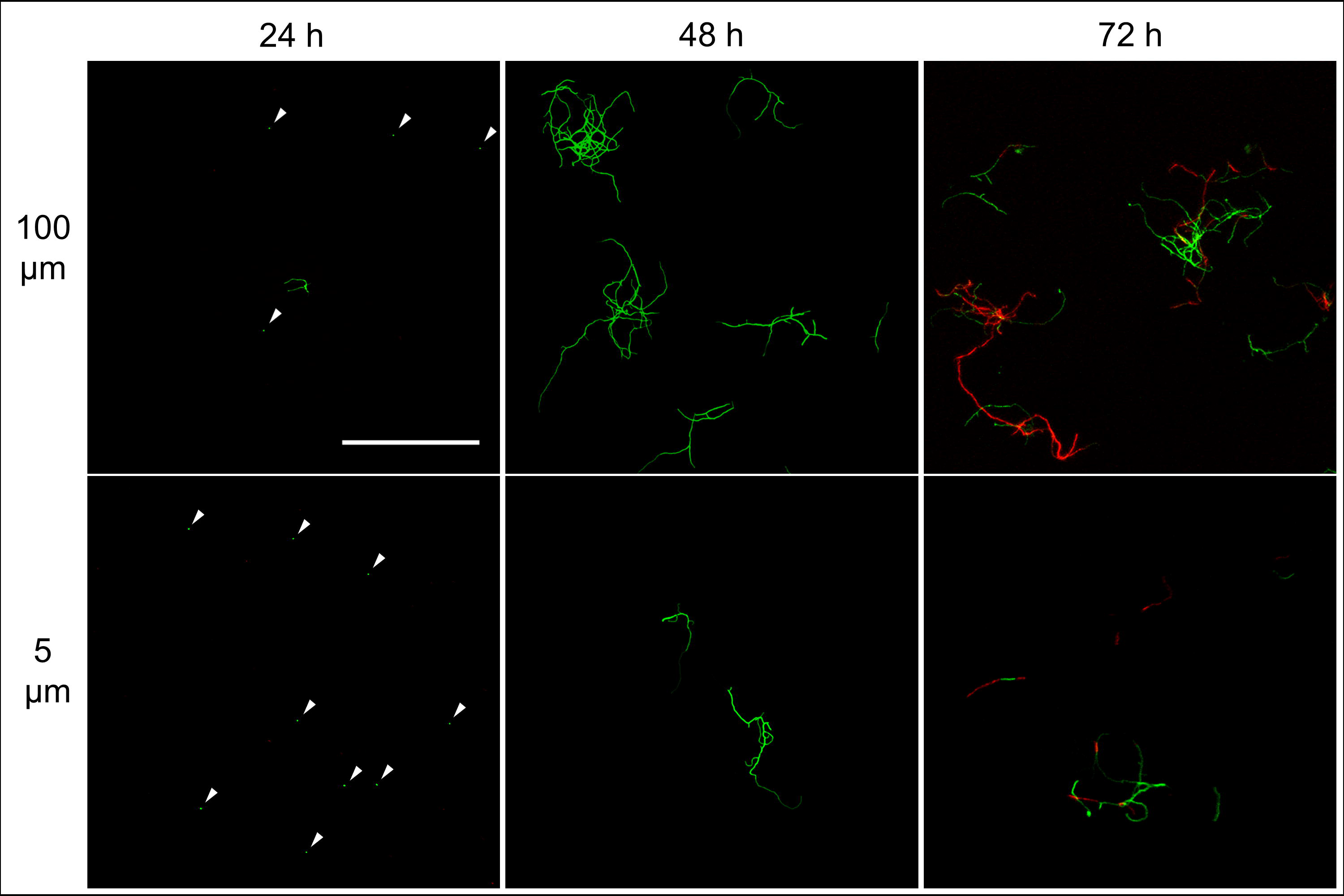
Morphological analysis of filtrates of seed cultures. Filtrates were obtained through the sequential filtering of TSBS cultures which had been grown for 24, 48 or 72 h, through cell strainers with a pore size of 100 (top panels), 40 and 5 μm (lower panels). Prior to imaging, samples were stained with Syto-9 (green) and propidium iodide (red) for the visualization of viable and dead mycelium, respectively. The scale bar represents 100 μm.

**Table 3.** Number of Colony Forming Units (CFU) in the filtrates of liquid-grown cultures of different ages. The used filters had an average pore size of 100 and 5 µm. The numbers were obtained by counting the Colony Forming Units (CFUs) after 48 hours of growth on MS agar medium.

| Culture age (h) | Pore size ($\mu\text{m}$) | CFUs $\text{ml}^{-1}$ |
| --- | --- | --- |
| 24 | 100 | $1,67 \pm 0.33 * 10^5$ |
| 24 | 5 | $1,63 \pm 0.51 * 10^5$ |
| 48 | 100 | $1,77 \pm 0.33 * 10^6$ |
| 48 | 5 | $1,62 \pm 0.53 * 10^6$ |
| 72 | 100 | $2.16 \pm 0.32 * 10^6$ |
| 72 | 5 | $1.91 \pm 0.28 * 10^6$ |

A markedly different situation was observed when the seed cultures were filtered after 48 h of growth. The number of CFUs in the filtrates had increased by an order of magnitude to 1.77*10^6^ ml^-1^ and 1.62*10^6^ ml^-1^ when using 100 and 5 µm filters, respectively (Table 3). Notably, the number of CFUs at this stage was higher than that of the inoculated spores, strongly implying that fragmentation had occurred. In agreement with this, a microscopic analysis of the filtrates revealed that small, viable mycelial fragments represented the vast majority of particles in the filtrate. After 72 h, filtered seed cultures also contained a large number of dead hyphae, in addition to viable fragments (Fig. 3).

### Fragmentation coincides with nutrient exhaustion in submerged cultures of *Streptomyces lividans*

To better analyze the occurrence of fragmentation throughout growth, we devised an experiment in which all particles smaller than 100 µm (spores and fragments) were first removed via filtration from seed cultures that had been grown for 16, 24, 32 or 40 h (see Material and Methods). The remaining particles were then brought back in the same medium and allowed to further grow for 8 h, after which the cultures were again filtered with a 100 µm filter. The number of viable particles in the filtrates was determined by counting the CFUs on solid MS agar plates. Using this method, we could quantify the occurrence of fragmentation within time frames of 8 h.

2,6*10^2^ CFUs ml^-1^ were found in the filtrate when 16-h old pellets were used. The number of CFUs increased more than 100-fold to 3.2*10^4^ and 7.0*10^4^ CFUs ml^-1^ with pellets of 32 and 40 h, respectively (Fig. 4, Table 4). Consistent with these observations, when the particles in the filtrates were brought into fresh TSBS medium and the biomass quantified that derived from their growth, a comparable trend was observed (Table 4).

**Figure 4.**
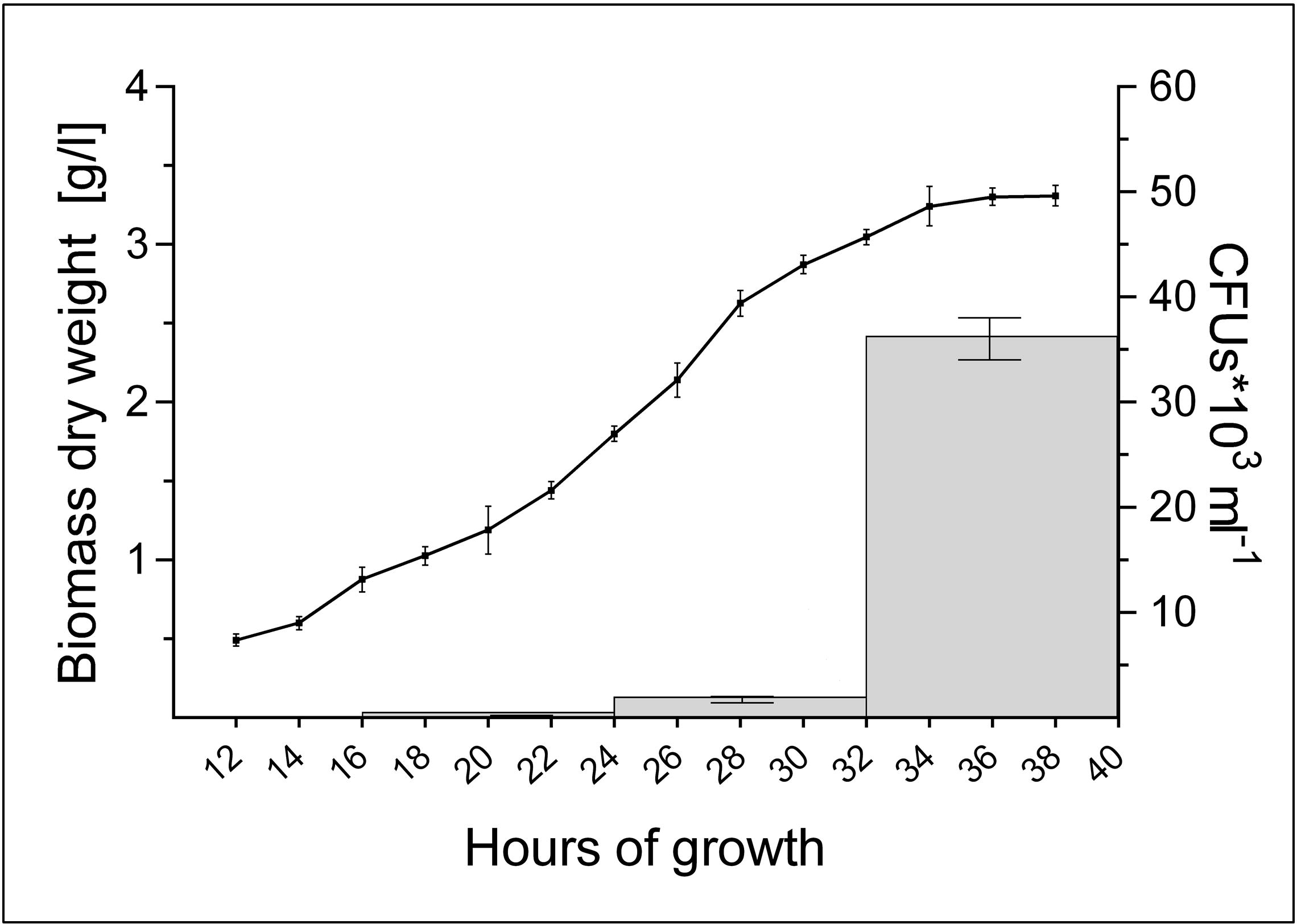
Abundant pellet fragmentation coincides with the entrance of cultures into the stationary phase. Growth curve of *Streptomyces lividans* 66 in TSBS medium, as determined by dry weight measurements over time (line). The grey bars represent the number of CFUs released by the growing pellets over time (see also Table 4).

**Table 4.** Number of viable fragments released in 8 hours time-frames by pellets collected from liquid-grown cultures of different ages. The numbers were obtained by counting the Colony Forming Units (CFUs) after 48 hours of growth on MS medium. The biomass (dry weight) obtained after inoculating the released fragments in fresh TSBS medium after 24 hours is indicated in g L^-1^.

| Culture age (h) | CFUs $\text{ml}^{-1}$ | Dry weight ($\text{g L}^{-1}$) |
| --- | --- | --- |
| 16 | $2.6 \pm 0.1 * 10^2$ | N.D. |
| 24 | $1.7 \pm 0.3 * 10^3$ | $0.3 \pm 0.1$ |
| 32 | $3.2 \pm 0.2 * 10^4$ | $3.1 \pm 0.2$ |
| 40 | $7.0 \pm 0.6 * 10^4$ | $3.7 \pm 0.2$ |

Notably, the steep increase in the number of viable particles in the filtrate correlated with the entrance of cultures into the stationary phase (Fig. 4). This prompted us to investigate whether the exhaustion of nutrients correlated directly with the onset of fragmentation. If true, we reasoned that fragmentation would occur earlier in cultures containing less nutrients. We therefore generated seed cultures of *S. lividans* in normal or half-strength TSBS (½ x TSBS) medium, which were used to inoculate fresh TSBS cultures after 12, 18, 24, 30 or 36 h. Fragmentation in half-strength TSBS medium occurred at least 12 h earlier than in normal TSBS medium, as inferred from the earlier detection of small pellets in the diluted cultures (see arrowheads in Fig. S3).

### Detached fragments are able to aggregate with one another, but not with mature pellets

In the previous section we have shown that the onset of fragmentation correlates with the entrance of cultures into the stationary phase. To test whether the detached fragments can aggregate with one another to form new composite pellets, we used fluorescent strains of *S. lividans* transformed with pGREEN (Zacchetti et al., 2016) and pRED*, constitutively expressing eGFP and mCherry, respectively. The seed cultures, containing equal amounts of each fluorescent strain, were diluted after 24, 48 or 72 h in fresh TSBS medium. Consistently with our previous results, we noticed that cultures inoculated with 24-h old seed cultures only contained large pellets, while many smaller pellets were found when 48- and 72-h old seed cultures were used (Fig. S4). Notably, all large and small pellets contained mycelium of both fluorescent strains. This suggests that the fragments contained hyphae of both fluorescent strains at the moment of detachment, or alternatively that detached hyphae expressing either fluorescent protein were able to aggregate to form composite pellets.

Visualization of the detached fragments collected from seed cultures of the co-cultured fluorescent strains indicated that they were invariably either green or red fluorescent (Fig. S5). This is consistent with a model in which the detached fragments are able to aggregate with one-another when forming new pellets. To test this more directly, cultures of the fluorescent *S. lividans* strains were grown first separately for 48 and 72 h. Equal amounts of each culture were subsequently transferred into fresh medium to give co-cultures and further grown for 24 h. Again, both large and small pellets occurred in these co-cultures. Fluorescence microscopy revealed that that 95.3 ± 3.8 % and 94.2 ± 3.1 % of the small pellets contained both red and green hyphae (Fig. 5) in the co-cultures performed with 48- and 72-h old seed cultures, respectively. Conversely, the large pellets were exclusively green or red fluorescent, showing that aggregation between small detached fragments and pre-existing mature pellets did not occur. To validate this observation, we repeated this experiment with a non-fluorescent wild-type strain and the strain constitutively expressing eGFP (Fig. S6), which would facilitate the detection of fluorescent hyphae associated with non-fluorescent pellets. However, no signs of adhesion between fragments and pre-existing pellets was observed. Altogether these results demonstrate that, when provided with fresh nutrients, the small detached fragments are able to aggregate with one another but not with pre-existing pellets.

**Figure 5.**
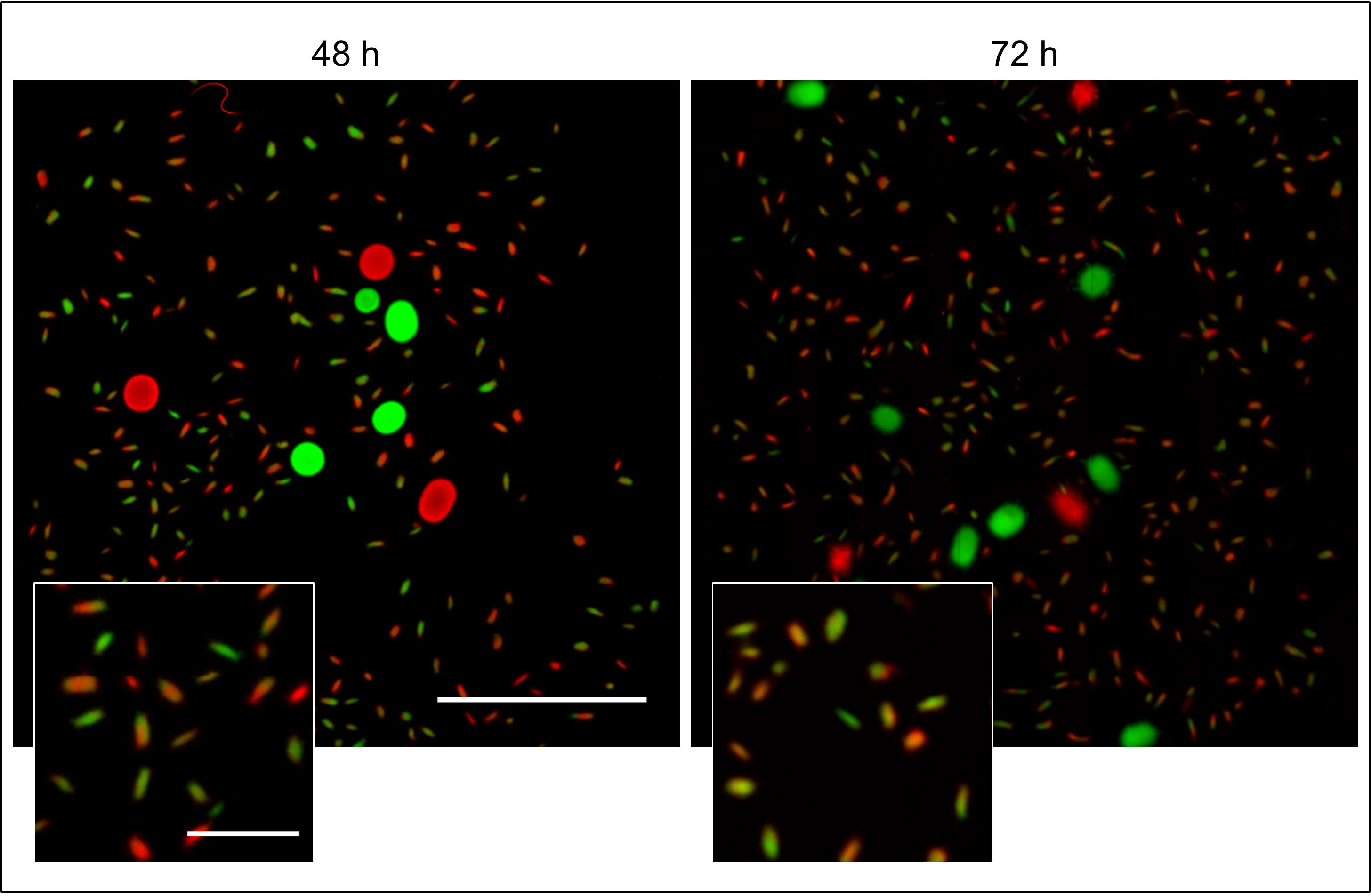
Detached fragments are able to aggregate with one-another during the formation of new pellets. Micrographs of pellets in co-cultures of red- and green-fluorescent derivatives of *Streptomyces lividans* 66 obtained by mixing separate cultures of these strains after 48 (left) and 72 h (right) of growth. The inlay shows that the small (new) pellets are composed of both red and green fluorescent mycelium. The scale bars represent 2 mm and 200 μm in the overview pictures and inlays, respectively.

### Shear stress limits aggregation between large particles

Studies addressing the impact of mechanical stress on the morphology of streptomycetes in liquid-grown cultures have demonstrated that high shear stress typically leads to the formation of small mycelial particles, while large pellets are observed with less shear (Tamura et al., 1997;Xia et al., 2014). Given that we only detected fragmentation after 30 h of growth, we reasoned that mechanical forces exert their influence on morphology largely by counteracting the aggregation between larger particles, rather than promoting fragmentation throughout growth. To test this hypothesis, we performed an experiment in which we separately grew the fluorescent derivatives of *S. lividans* for 12 h, when the aggregation between distinct particles is known to become negligible (Zacchetti et al., 2016). We then mixed these cultures in flasks either with or without a metal coil, a stratagem commonly used to regulate the degree of shear stress (Kieser et al., 2000). Consistently with earlier observations, the presence of the metal coil prevented aggregation between pre-existing particles, as concluded from the observation that only 3.7 ± 1.0 % of pellets were either red or green fluorescent (Fig. 6). In co-cultures performed without coils, however, we found that 68.7 ± 0.4 % of pellets contained distinct patches of red and green fluorescent mycelium. On average, these composite pellets were larger (352.3 ± 87.7 µm) than those formed in higher shear stress conditions (179.8 ± 37.4 µm). Taken together, these results indicate that mechanical stress plays a role in shaping pellets by limiting the aggregation between large aggregates.

**Figure 6.**
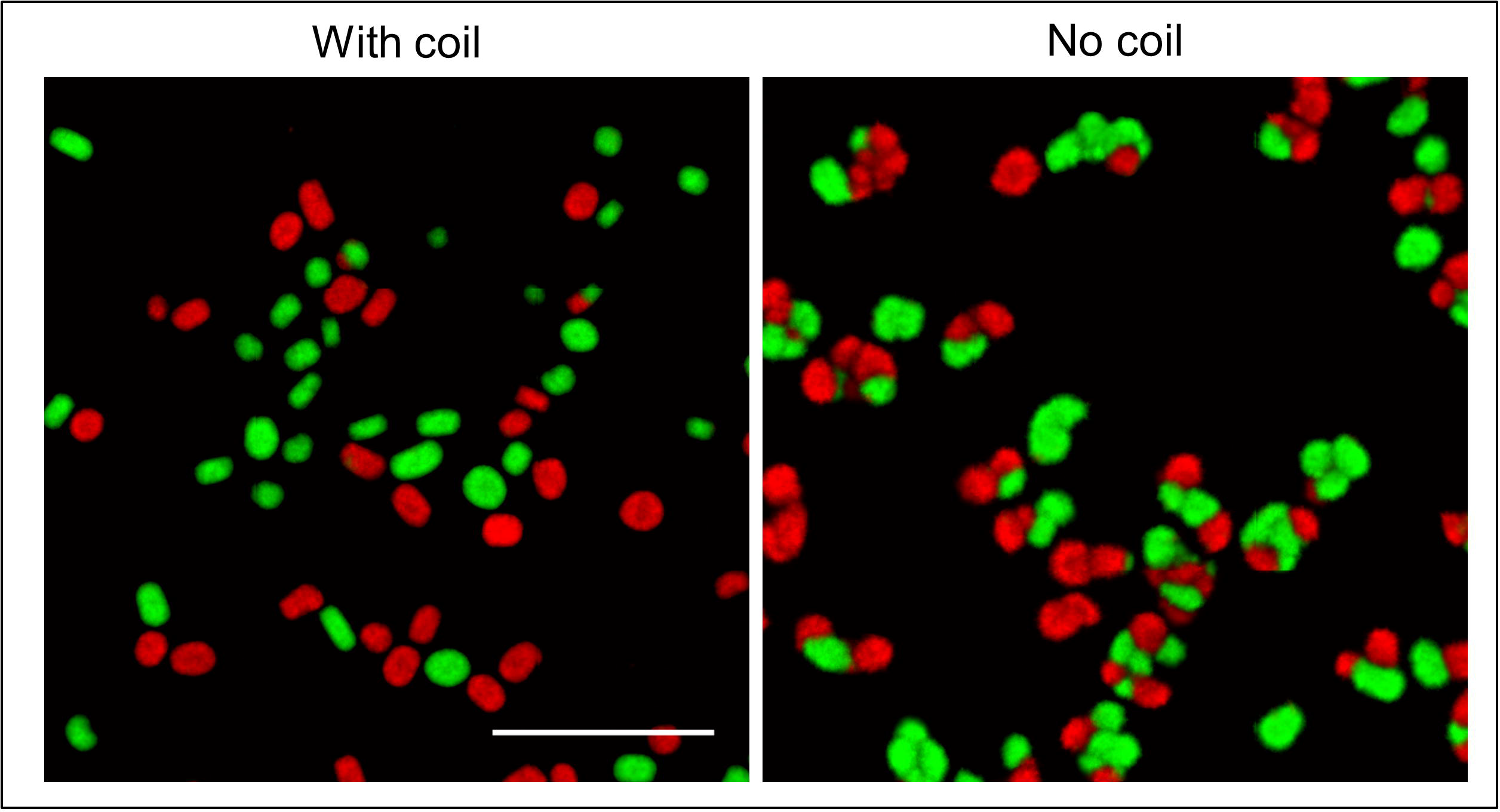
Shear stress limits aggregation between large particles: Micrographs of pellets in co-cultures of the red- and green-fluorescent derivatives of *Streptomyces lividans* 66. These co-cultures were obtained by mixing separate cultures of the fluorescent strains after 12 h of growth and allowing these to grow for 24 h in flasks with (left) or without (right) a shear-inducing coil. The scale bar represents 1 mm.

## Discussion

Over the past years we have witnessed a dramatic increase in our understanding of the physiology and growth dynamics of liquid-grown streptomycetes (van Dissel et al., 2014). Dissecting the factors that influence the submerged morphology of these bacteria is of crucial importance to improve their industrial exploitation, given the correlation between morphology and production (Wardell et al., 2002;van Wezel et al., 2006). We here performed a systematic analysis of fragmentation in submerged cultures of the industrial workhorse *S. lividans* and show that this process is initiated when cultures enter a stationary phase. Notably, when fresh nutrients are provided, the released fragments are able to establish new pellets. This behavior is strikingly similar to the dispersal of biofilms, a strategy that allows the inhabitants of these multicellular communities to colonize other niches when the environmental conditions become unfavorable (e.g. nutrient starvation, presence of toxins).

The similarities between *Streptomyces* pellets and biofilms are remarkable also from a broader perspective (Fig. 7), as previously suggested (Kim and Kim, 2004). As in biofilms, the formation of pellets depends on the ability to synthesize extracellular polymeric substances (EPSs; (Flemming et al., 2007;Flemming and Wingender, 2010). *S. lividans* produces at least two EPSs, which are a glycan containing β-(1-4)-glycosidic bonds produced under control of the *cslA-glxA* operon, and PNAG, which is synthesized by the MatAB proteins (Chaplin et al., 2015;van Dissel et al., 2015;van Dissel et al., 2018). PNAG is a widespread component of bacterial biofilms, e.g. those formed by *Staphylococcus*, but also many other species (Götz, 2002;Jefferson, 2009). In *Streptomyces*, these glycans mediate the adhesion between hyphae either belonging to the same or to distinct particles, a feature that causes the deletion mutants of *cslA*, *glxA or matAB* to grow as individual particles with an open morphology (Zacchetti et al., 2016).

**Figure 7.**
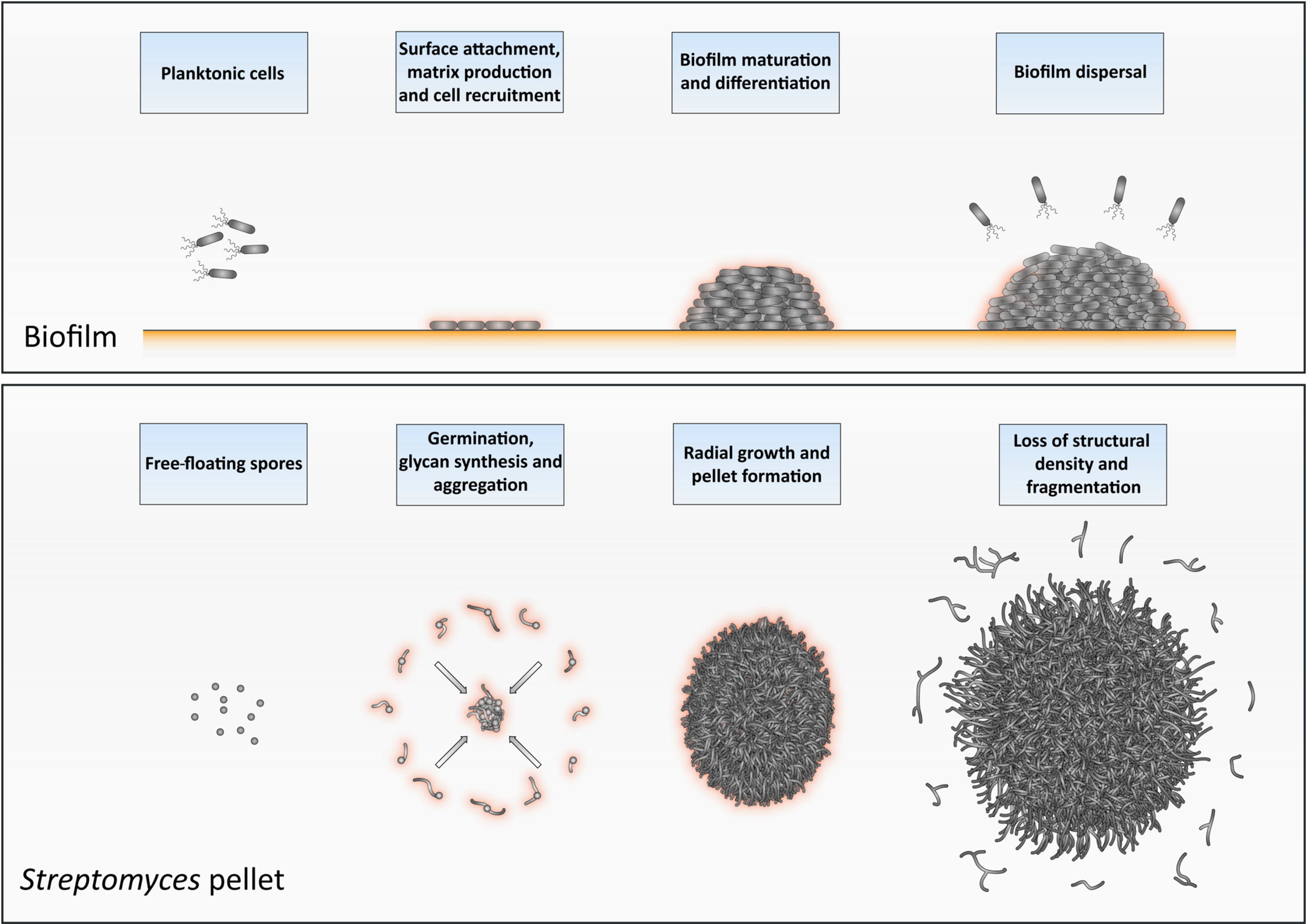
Similarities between the formation and development of biofilms and of *Streptomyces* pellets. At the onset of biofilm formation, planktonic cells differentiate into non-motile, matrix-producing cells that attach to a surface and recruit other cells by means of their adhesiveness. This leads to the formation of a large community of cells that are embedded in the extracellular matrix. When the conditions become unfavorable, such as when nutrients and other resources become limiting and waste products accumulate, the process of biofilm disassembly is initiated. In this way, cells are released from the biofilm microstructure to disperse to more favorable locations. The growth cycle of *Streptomyces* pellets begins with the germination of spores. The adhesive glycans that cover the surface of the so-called germlings cause them to rapidly adhere to one-another and to other small aggregates. The aggregates further expand radially utilizing the available nutrients to eventually form dense pellets. When the stationary phase of growth is reached, the outer edge of pellets becomes fluffier due to the appearance of protruding hyphae, which detach as fragments. When provided with fresh nutrients, these fragments are able to establish new pellets.

In *S. lividans*, the aggregation of distinct individuals occurs during the early stages of growth. This process has a duration that depends on the degree of mechanical stress in liquid cultures. Here we show that when the level of shear is reduced, larger particles are able to further aggregate, after which they continue to grow as a larger whole. When a threshold size is reached by these particles that impedes further aggregation, they further increase in size as a result of hyphal elongation and eventually form pellets. Pellets may also become fortified at this stage by other extracellular polymers, including amyloid fibrils, extracellular DNA and hyaluronic acid (Kim and Kim, 2004). The involvement of these components in strengthening the adhesion between cells is commonly observed during the development and maturation of various bacterial biofilms (Flemming and Wingender, 2010;Abee et al., 2011).

Opposite to pellet formation is the process of their disassembly, commonly referred to as fragmentation. In this study, we provide compelling evidence that fragmentation only commences when cultures enter their stationary phase. From this moment onwards, small mycelial fragments are found in the culture broth and their number increases rapidly over time. Initially, the majority of these fragments are viable, able to aggregate with one another and to form new pellets when provided with fresh nutrients. In the absence of fresh nutrients, however, these fragments eventually die. Interestingly, the onset of fragmentation coincides with a change in the morphology of pellets. More specifically, numerous hyphae protrude from the periphery of aging pellets (see Fig. 2 and S2). The adhesive forces between hyphae may be weakened at this stage, making these filaments prone to fragmentation as a result of mechanical stress. Given that fragmentation is initiated when cultures enter the stationary phase, we speculate that this process may also be stimulated by the organism itself, similarly to the ordered process of biofilm disassembly (Kaplan, 2010;McDougald et al., 2011;Guilhen et al., 2017). Processes related to programmed cell death (PCD), which are known to occur in aging *Streptomyces* pellets, could contribute to the active detachment of fragments (Manteca et al., 2008;Rioseras et al., 2013). Furthermore, several organisms employ specific hydrolases to degrade the endogenous extracellular matrix and promote the dispersal of cells. An interesting example to mention in this context is the PNAG-hydrolase Dispersin B, produced by *Actinobacillus actinomycetemcomitans* (Kaplan et al., 2003). Dispersin B is also able to counteract the deposition of PNAG on the hyphal surface of *Streptomyces* pellets (van Dissel et al., 2018). Interestingly, a putative homologue of dispersin B is present in *S. lividans*. It thus seems conceivable that pellet-forming streptomycetes could also actively contribute to the disassembly of pellets and the release of fragments as a strategy to maximize the chances of survival in hostile environments.

Altogether, our work provides new valuable insights into the dynamics of pellet formation and disassembly in the industrial workhorse *S. lividans*. Besides the fundamental interest, we envision that these results are useful for a better understanding and more rational optimization of production processes. In particular, the notion that inoculating mycelium of different ages results in differently sized populations of pellets can be very important for the accumulation of biomass in industrial processes, especially when a particular morphology is preferred for the production of the desired product.

## Acknowlegments

We thank Erik Vijgenboom, Gilles van Wezel and Dino van Dissel for fruitful discussions. This work was supported by a VIDI grant (12957) from the Dutch Applied Research Council to DC.

## Author Contributions

BZ and DC designed the experiments, which were performed by BZ and PS. The manuscript was written by BZ and DC, with input from PS.

## Conflict of Interest Statement

The authors declare that the research was conducted in the absence of any personal, professional or financial relationships that could potentially be construed as a conflict of interest.

---

**Figure S1: Size distributions of pellets in diluted cultures of *Streptomyces lividans* 66**. TSBS cultures were inoculated with 2 or 10 ml of seed cultures that had been grown for 24, 48 or 72 h. The plots represent the size distribution of at least 300 pellets per sample, obtained after 24 h of growth. All sizes are indicated in micrometers.

**Figure S2. Morphological changes accompanying the aging of *Streptomyces* pellets.** Collage of representative micrographs of pellets of *Streptomyces lividans* 66 in TSBS cultures after 24 (top) and 72 h (bottom) of growth. The scale bars represent 200 μm.

**Figure S3. Fragmentation correlates with nutrient availability.** Micrographs representing pellets of *Streptomyces lividans* 66 obtained by diluting seed cultures in fresh TSBS medium after 12, 18, 24, 30 and 36 h of growth. The seed cultures were prepared in normal TSBS (top panels) or in ½ x TSBS (bottom panels) medium. Note that the small pellets, derived from outgrowing fragments, are observed at least 12 h earlier in ½ x TSBS than in normal TSBS. The scale bar represents 1 mm.

**Figure S4. Morphological analysis of pellets of the fluorescent derivative strains of *Streptomyces lividans* in TSBS cultures.** Seed cultures were prepared by co-culturing *S. lividans* strains constitutively expressing eGFP or mCherry in TSBS medium. The morphology of pellets in these seed cultures after 24 (left), 48 (middle) and 72 (right) h of growth are shown in the top panels. The bottom panels show micrographs of pellets following the transfer of 10 ml of the seed cultures in fresh TSBS medium and subsequent growth for 24 h. The scale bar represents 1 mm.

**Figure S5. Visualization of detached fragments from pellets of co-cultured fluorescent *S. lividans* strains.** Filtrates were obtained by the sequential filtering of TSBS cultures, which had been grown for 48 or 72 h, through cell strainers with a pore size of 100 (top panels), 40 and 5 μm (lower panels). Note that the detached fragments are either green or red fluorescent. The scale bar represents 100 μm.

**Figures S6. Large, fragmenting particles are inert to aggregation**. Micrographs of pellets from co-cultures of the *S. lividans* wild-type strain and its green-fluorescent derivative, obtained by mixing separate cultures of both strains after 48 (left) and 72 h (right) of growth. The inlay shows that the large wild-type pellets remain non-fluorescent after the transfer, indicating that small mycelial fragments do not aggregate with these large particles. The scale bars represent 2 mm and 200 μm in the overview pictures and inlays, respectively.

